# The Effects of Copper Addition on The Structure and Antibacterial Properties of Biomedical Glasses

**DOI:** 10.1101/2020.01.24.918524

**Authors:** Leyla Mojtabavi, Amir Razavi

**Affiliations:** University of Tehran, Department of Engineering Science, Enghelab Ave, Tehran, Iran 111554563

**Keywords:** Bioglass, copper, structure, XPS, antibacterial

## Abstract

In this work, we studied the effects of copper incorporation in the composition of bioactive glass. Three different glass compositions were synthesized with 0, 3, and 6 mol% of copper addition. X-Ray Diffraction (XRD) patterns confirmed that an amorphous microstructure was obtained for all three glass compositions. Results from Differential Thermal Analysis (DTA) showed that the copper addition in the glass lowers the glass transition temperature, from 646°C to 590°C when added at 6 mol%. X-ray Photoelectron (XPS) survey and high-resolution scans were performed to study the structural effects of copper addition in the glass. Results indicated that the incorporation of copper changes the ratio of bridging to non-birding oxygens in the structure. Glasses were further analyzed for their structure with Nuclear Magnetic Resonance (NMR) spectroscopy, which indicated that copper acts as a network modifier in the glass composition and copper-containing glasses show a less connected microstructure. Antibacterial efficacy of the glasses was analyzed against E. coli and S. epidermis. Copper-containing glasses showed a significantly higher inhibition zone compared to control glass. The glass with 6 mol% copper, exhibited inhibition zones of 9 and 16mm against E. coli and S. epidermis bacteria, respectively.

## Introduction

Bioglass®, also known as 45S5 glass, was first discovered by professor Larry Hench in the late 1960s in order to develop a material to fill bone voids without being rejected by the body^1^. The initial 45S5 Bioglass® that was introduced by Hench was a silicate-based glass with a composition of 45SiO_2_-24.5Na_2_O-24.5CaO-6P_2_O_5_ in weight percent^2^. Early studies on this composition showed its capacity to form a chemical bond with living tissue^3^. As a biodegradable material, the bioglass degrades slowly in an aqueous solution and stimulates new bone formation and growth through a chemical bond to the existing bone tissue^4^. The dissolution of the glass and formation of the calcium phosphate layer on the surface of glass initiates the osteogenesis mechanism. In addition, the release of dissolution products from the glass into surrounding physiological solutions can provide therapeutic effects to adjacent tissue^5^. In the past few years, different compositions of bioactive glasses have been developed to improve the properties and take full advantage of these glasses for biomedical applications. The bioactive glasses have been used in dental applications, applied as bioactive coatings on the surface of metal implants, used as bioactive components of composites and have been made into synthetic bone scaffolds^2-3, 6-8^. The diversity of their applications comes from the glass composition, and how it can be modified by the incorporation of new elements and therefore changing the glass structure^2^.

The atomic structure of the glass significantly affects its properties and in particular the bioactivity and degradation rate which directly influences the formation of calcium phosphate surface layer and cell viability^9^. The key compositional factor in the determination of the bioactivity of these glasses is the SiO_2_ content^10^. The bioactive glass composition consists of relatively low amounts of silica and high concentrations of network modifiers such as Ca and Na to facilitate its dissolution in the physiological solutions^8, 10^. The structure of bioactive glasses can be described in the same way as other silicate glasses which have been well characterized in terms of the arrangement of the SiO_4_ tetrahedron coordination^11^. The number of tetrahedrons determines the size of the ring used to characterize the network. The structure of pure SiO_2_ consists of a three-dimensional network of SiO_4_ tetrahedra, and each tetrahedron is associated with bridging oxygen in the structure, through Si-O-Si bonding^12^. The addition of network modifying elements disrupt the linkage of SiO_4_ tetrahedra and creates non-bridging oxygens^13^. As a result of modifier ions incorporation, the disrupted glass network is now prone to hydrolysis in aqueous solutions, which is a critical step for the bioactivity of glass^5^.

Since the introduction of bioactive glass by Hench, a wide variety of chemical modifications have been proposed by the incorporation of different elements in the glass. Some very common elements are Zn, Ti, and Sr^14-15^. These elements have been added to the glass composition to improve the glass physical, mechanical and biological behavior^9, 16^. In this work, we have synthesized copper incorporated bioactive glass composition. Copper is a monovalent/divalent cation acting as a network-modifier in the glass structure^17^. The addition of CuO at the expense of SiO_2_ can make the glass composition more soluble in the physiological solution, which in turn can enhance the bioactivity. Irrespective of its role, copper may be beneficial to the biological properties associated with the resultant material as there are numerous reports on the positive antibacterial response associated with the use of copper^18-19^. The main purpose of this study is to modify the glass composition with the incorporation of copper in the glass and to determine its structural effects on the glass network. Further, we aim to investigate any changes occurring in the antibacterial efficacy of the glass as a result of copper addition.

## Materials & Methods

### Glass Synthesis

Three different glass powders were synthesized; a control and two copper-containing glasses. The control glass (*BG*) was synthesized based on a SiO_2_-CaO-Na_2_O-P_2_O_5_ composition. The copper-containing glasses contain 3 and 6 mol% of CuO at the expense of SiO_2_ (Table 1). The analytical grade reagents of SiO_2_, CuO, CaCO_3_, Na_2_CO_3_, NH_4_H_2_PO_4_ were mixed and milled for 30 minutes. The glass batch was then dried at 110°C for 1 hour and melted at 1400°C for 5 hours in a platinum crucible and shock quenched into water.

**Table 1.** Glass compositions (mol%) where SiO_2_ is substituted with CuO.

|  | SiO<sub>2</sub> | CaO | Na<sub>2</sub>O | P<sub>2</sub>O<sub>5</sub> | CuO |
| --- | --- | --- | --- | --- | --- |
| *BG* | 55 | 20 | 20 | 5 | 0 |
| *BG-3Cu* | 52 | 20 | 20 | 5 | 3 |
| *BG-6Cu* | 49 | 20 | 20 | 5 | 6 |

### X-Ray Diffraction

Synthesized glass powders were analyzed for their diffraction patterns using X-Ray Diffractometer (Shimadzu XRD-6100X) at 40 kV and 30 mA utilizing CuKa radiation. The range of angles was from 10° to 60°, at a step size of 0.03° and step time of 10 s.

### Differential Thermal Analysis

Differential Thermal Analysis (DTA) experiments were performed using a SDT Q600 DTA instrument. Around 25 mg of each of the glass powders were loaded in platinum pans, where one empty platinum pan was also used as a reference. DTA curves were collected in air from 25°C to 1200°C at the heating rate of 10°C/min.

### X-Ray Photoelectron Spectroscopy

X-Ray Photoelectron Spectroscopy (XPS) was performed using a VG-Microtech ESCA-2000 system with AlKa source (hν=1486.6 eV), at output of 25.5 W. Before the sample loading, the glass powders were first kept in a vacuum oven at 90°C for 30 min to allow drying of the powders. During the test, the vacuum in the analysis chamber of instrument was in the range of 10^−9^–10^−10^ mBar. Survey scans, and high-resolution spectra of glass powders were collected, and the binding energies referenced to the C1s at 284.6 eV.

### Magic Angle Spinning Nuclear Magnetic Resonance (MAS-NMR)

MAS-NMR was carried out on powdered glass samples using a DSX-200 FT-NMR. The measurements were conducted at a frequency of 39.77MHz, with the spinning rate at the magic angle of 5 kHz for 29Si with a recycle time of 2.0s. The reference material used to measure the chemical shift was tetramethylsilane (TMS).

### Antibacterial Agar Diffusion

The antibacterial efficacy of the glasses was tested using the agar diffusion test method. First, glass disks of the synthesized powders were prepared and kept under UV for 4 hours for sterilization. Bacteria was grown aerobically in liquid broth at 37°C for 48 hours. Glass disks were placed in petri-dish covered with bacterial agar. The agar plates were then inoculated with the bacteria solution using a sterile swab, and kept in the incubator for 24 hours at 37°C. After 24 hours passed, the samples were then evaluated for their inhibition zone by measuring the size of the zone from the edge of the disk to the end of inhibition zone.

## Results & Discussion

In this study, a series of bioactive glass compositions were synthesized. *BG* is a control bioactive glass composed of SiO_2_, CaO, Na_2_O, and P_2_O_5_. *BG-3Cu*, and *BG-6Cu* glasses contain incremental amounts of 3 and 6 mol% of CuO at the expense of SiO_2_. The specific compositions are shown on Table 1. The initial characterization on glasses was to generate X-ray diffraction patterns to identify if any crystalline phases were present in the glasses. XRD was performed on each glass composition (*BG, BG-3Cu*, and *BG-6Cu*), and results are presented in Figure. 1. All three glasses, show the characteristic amorphous curve for the silicate glasses at around 20 to 30 degrees. The amorphous hump arises from the glasses having no long-range atomic order in their molecular arrangement. The results from the glasses diffractograms shows that the glasses used for this study were predominantly amorphous in structure.

**Figure 1.**
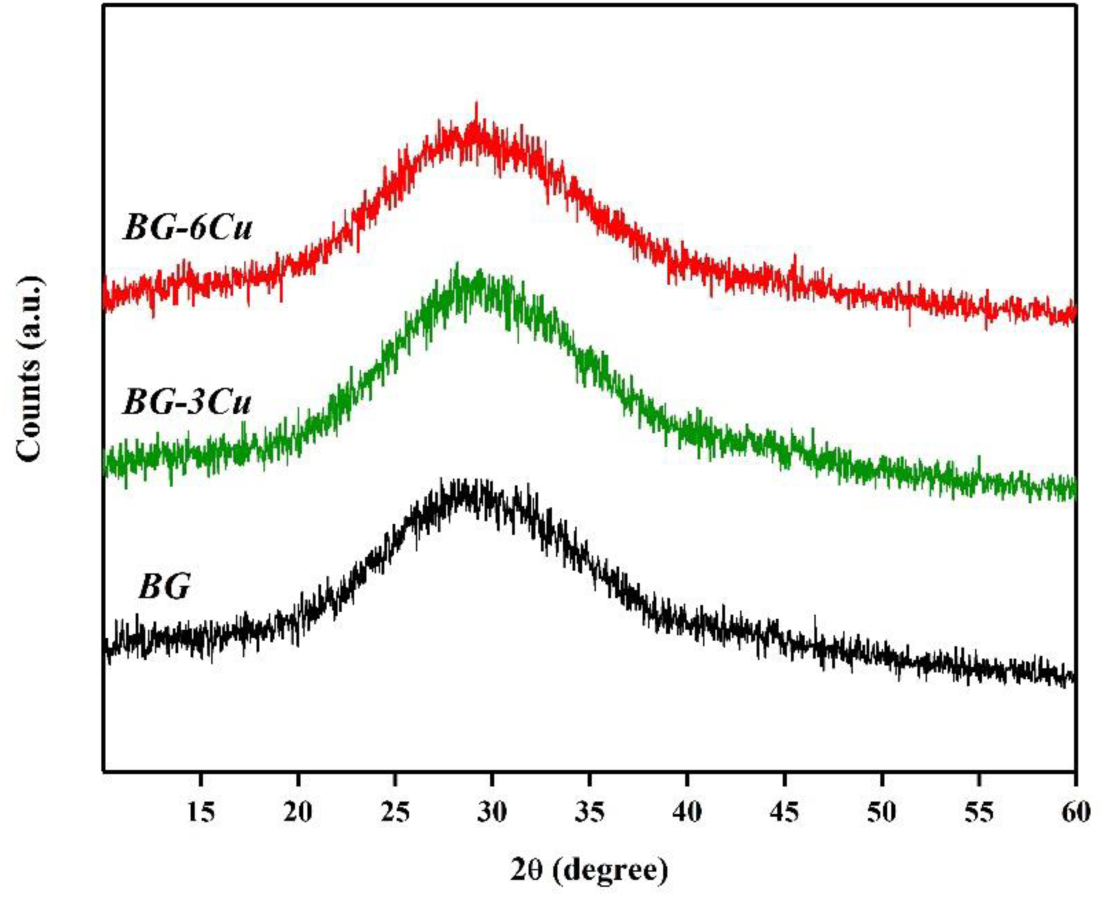
X-Ray diffractograms of amorphous glass powders of *BG, BG-3Cu*, and *BG-6Cu*.

Differential thermal analysis (DTA) was employed to measure the glass transition temperature (Tg), and the glass crystallization temperature (Tc), as a function of CuO addition. Figure 2 shows the DTA profiles of the glass series. The glass transition temperature (Tg) of BG was found to be 646°C, and the only crystallization peak (Tc) started at 936°C. The Tg of *BG-3Cu*, and *BG-6Cu* glasses were found to be at 611°C and 590°C, respectively. *BG-3Cu* showed two separate crystallization exotherm starting at 752°C and 864°C. However, *BG-6Cu*, only had one crystallization exotherm starting at 901°C. From Figure 2 it can be seen that the CuO incorporation in the glass chemistry, shifts the glass transition to lower temperatures. Tg represents the temperature at which the glass begins transformation from a solid material to a supercooled viscous liquid. Below Tg the bonding in amorphous glass particles remains in a solid state, however as the temperature increases the bonding within the glass becomes more mobile where the materials behave more like a liquid. Tg is an important materials property as it represents the upper temperature limit at which the glass can be used. In this case, Tg establishes the network disruption of the glass, where a more disrupted glass network has a lower Tg. Therefore, DTA is a useful technique, which can be used to determine if any differences in the glass structure exists between the glasses as a function of temperature. Preliminary analysis of the glass structure using DTA, suggests that the CuO addition decreases the network connectivity and hence increases its disruption. This further investigated by XPS and NMR studies.

**Figure 2.**
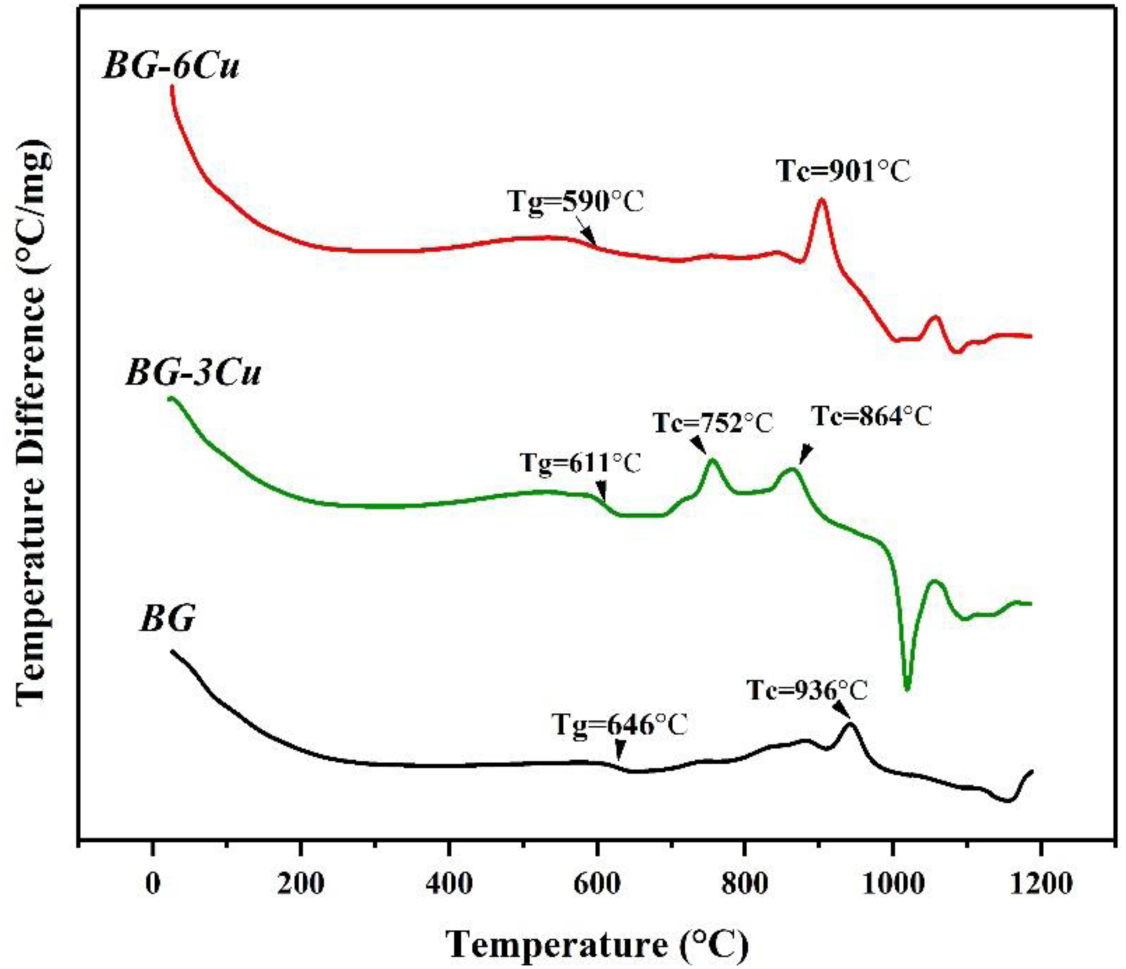
DTA profiles of *BG, BG-3Cu*, and *BG-6Cu* glasses. Glass transition, and crystallization temperatures are marked on each thermogram.

Compositional analysis of each glass was then determined using X-ray photoelectron spectroscopy (XPS) survey scans as shown in Figure 3. The survey scan of *BG* contains Na, O, Ca, P, and Si with minor traces of carbon (C). Both *BG-3Cu*, and *BG-6Cu* glasses show presence of Na, O, Cu, Ca, P, and Si with minor traces of carbon (C) as well, which corresponds with initial glass batch composition as shown in Table 1. X-ray photoelectron spectroscopy (XPS) survey scans were used to confirm the glass compositions and to find any possible contamination within the samples. High-resolution X-ray photoelectron spectroscopy (XPS) was conducted on glasses to study the structure of glass as a function of CuO addition. The XPS O1s signals of glass series are shown in Figure 4. There is a compositional dependence on the binding energy of the O1s spectra of glasses when CuO is introduced in the composition. The peak for the control *BG* glass is centered at 532.93 eV. For the copper-containing glasses, the O1s binding energies were found to be at 532.71, and 532.59 eV for *BG-3Cu*, and *BG-6Cu*, respectively. As can be seen from Figure 4, the incorporation of the Cu at the expense of Si shifts the O1s binding energy to lower values. Within the structure of oxide glasses, different oxygen species are present, as oxygen ions in the glass have different types of chemical bonding^20^. The bonding environment of the oxygen ions can be studied by analyzing the O1s spectrum of the glass. Therefore, XPS O1s is a very useful technique to study the contribution of bridging to non-bridging oxygens in the glass^21^. The lower binding energy peak of the O1s spectra is attributed to non-bridging oxygens, whereas the higher binding energy peak represents the bridging oxygens. The shift of the binding energies to lower values within the structure of copper glasses indicate an increase in the number of non-bridging oxygen at the expense of bridging oxygens^21^. This effect is suggesting that the Cu is acting as a network modifier in the glass by breaking the Si-O-Si bonds and replacing them with Si-O-NBO. This is in agreement with the glass transition temperature studies observed with DTA. Glasses were further analyzed for their structure with ^29^Si MAS-NMR.

**Figure 3.**
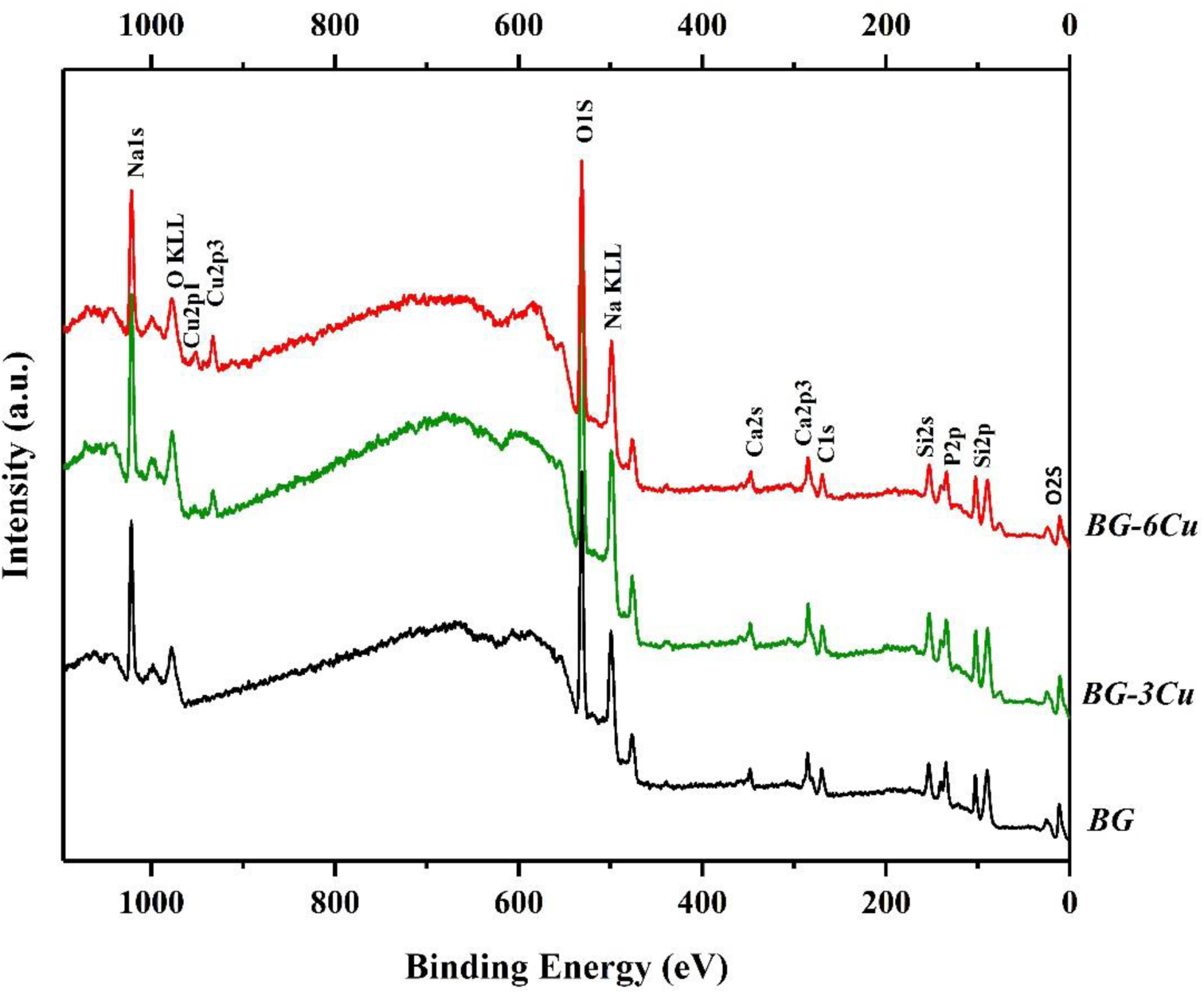
XPS survey scans of *BG, BG-3Cu*, and *BG-6Cu* glasses.

**Figure 4.**
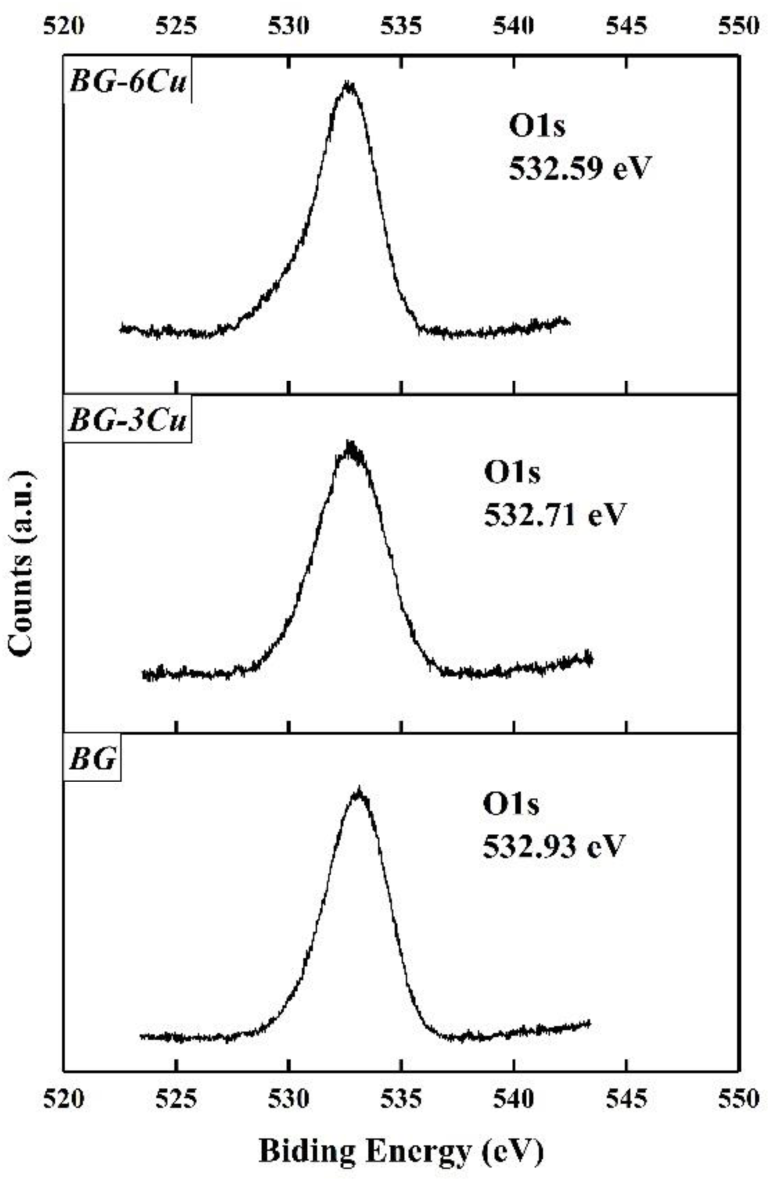
XPS O1s spectra of *BG, BG-3Cu*, and *BG-6Cu* glasses.

Magic Angle Spinning Nuclear Magnetic Resonance Spectroscopy (MAS-NMR) was employed to analyze the structure of the glass series. Figure 5 shows the ^29^Si MAS-NMR spectra of the glass series, as SiO_2_ is the network former in the glass composition. The chemical shift indicates the local environment around the silicon atom. The silicon spectra in these glasses predominantly show a large broad peak at around −75 to −105 ppm. The chemical shift in a four-coordinate silicon state has been found to be between –60 and –120 ppm^22^. The chemical shift of the spectra in the glass depends on the percentage of non-bridging, and bridging oxygens bound to the silicon. Increasing the number of non-bridging oxygens will increase the shielding of the nucleus of the atom, so the peak shifts to the positive direction. However, increasing the number of bridging oxygens moves the peak in a negative direction due to decreased shielding^23^. The Q-structure of the glass relates to the number of BOs associated with the silicon atom. A silicon atom with 4-BOs can be assigned a Q-structure of Q^4^, 3-BOs assumes a Q^3^-structure, 2-BOs assumes a Q^2^-structure etc^24^. MAS-NMR can be used to calculate the chemical shift position for identifying the Q-structure of silica glass. Chemical shifts have been identified at >-78ppm for Q^1^, −80 ppm for Q^2^, −90ppm for Q^3^ and −110ppm for Q^4^ ^22^. The MAS-NMR spectra of the glasses tested in this work, indicates predominant Q^3^/Q^2^ structures due to the chemical shift occurring mostly between −85 and −95 ppm. However, the broadness of the peaks suggests that all Q-structures may be present in the glass. For Control glass (*BG)*, a high degree of asymmetry exists within the Q^4^ region of the spectra. The peaks obtained from the MAS-NMR spectra of the *BG, BG-3Cu*, and *BG-6Cu* glasses are centered at −91 ppm, −89 ppm, and −88 ppm, respectively. It is evident with copper-containing glasses, that their broad spectra predominantly occur at Q^3^ structure, however, a slight shift is happening towards the Q^2^ region of the spectra as they move more towards the positive direction. This suggests that copper glasses may contain higher quantities of Q^2^ structures and therefore contain higher number of non-bridging oxygens compared to control. The results obtained from DTA, XPS and NMR spectra, suggest that the control glass is more structurally defined than the copper-containing glasses. Incorporation of the copper in the glass disrupts the glass network by increasing the number of non-bridging oxygens, which indicates that CuO acting as a network modifier in this glass system. Understanding the distribution of Q-species is important because the formation of non-bridging oxygens will have an effect on glass solubility, which plays an important role in dissolution of the particle surface that initiates the bioactive sequence^25^.

**Figure 5.**
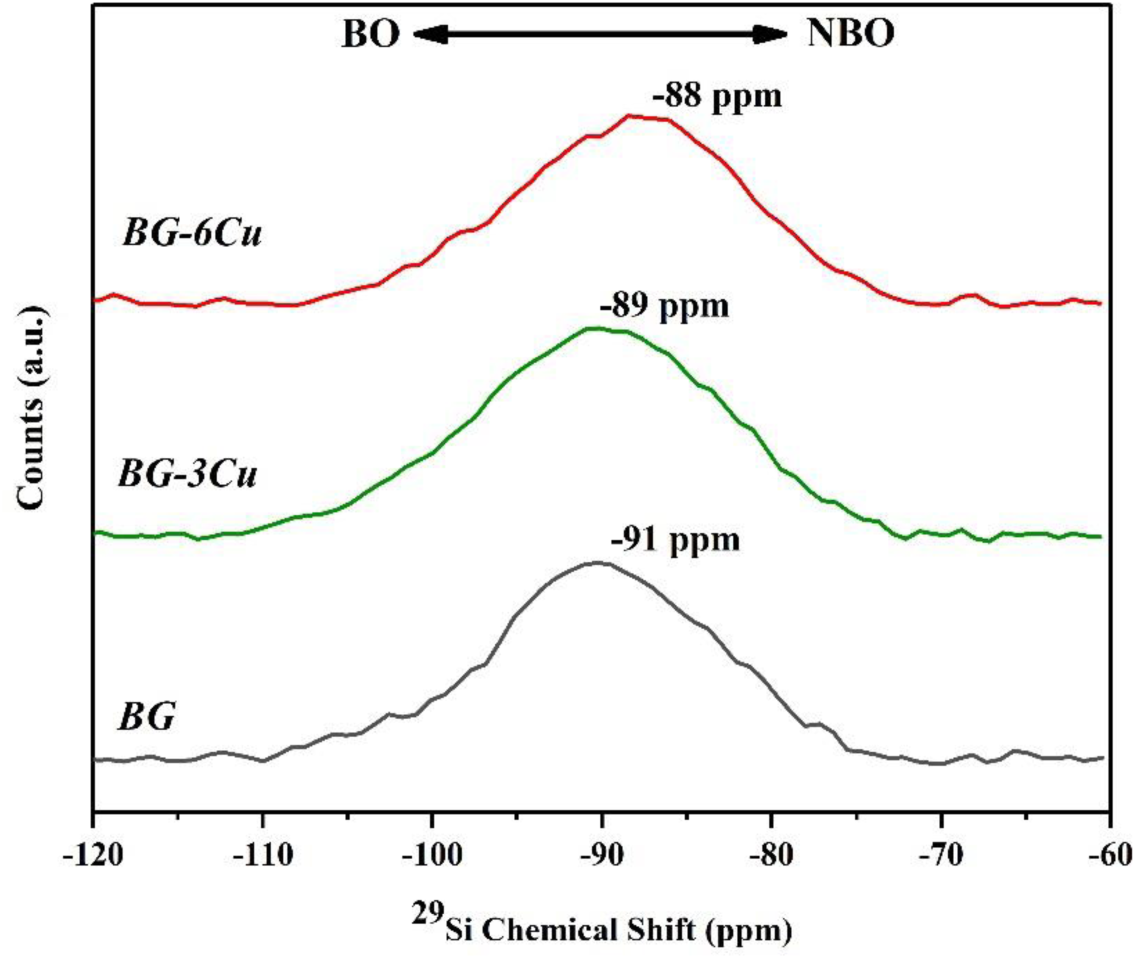
^29^Si MAS-NMR spectra of glass series.

One of the primary advantages of bioactive glass compositions is the ability to incorporate therapeutic ions to improve the antibacterial efficacy of the glass and therefore to reduce the infection risk^16^. The antibacterial efficacy of the glass compositions was evaluated using agar diffusion testing method and against two common bacteria strains: aerobic Gram-positive coccus S. epidermis and aerobic Gram-negative, E. coli. Glass disks were placed within inoculated plated, cultured for 24 hours, and then tested for their inhibition zones. The results from the agar diffusion test of the glass samples are presented in Figure 6. The copper-containing glasses are showing significantly higher antibacterial efficacy compared to control glass. No inhibition zone was detected for control glass against E. coli, however for *BG-3Cu*, and *BG-6Cu* glasses, 3 and 9mm of inhibition zone was observed. Glasses showed better antibacterial properties against S. epidermis bacteria, with 2, 5 and 16 mm of inhibition zone for *BG, BG-3Cu*, and *BG-6Cu* glasses, respectively. From Figure 6, it is evident that the inhibition zones associated with the copper glasses increases with the copper concentration in the glass. The antibacterial mechanism of copper can be complex, however previous studies suggest that copper ions disrupt the function of microbes by binding to proteins and enzymes in the microbes and therefore impede their activity. Using copper in the composition of the bioactive glasses can provide an alternative to antibiotics loaded biomaterials^18, 26^.

**Figure 6.**
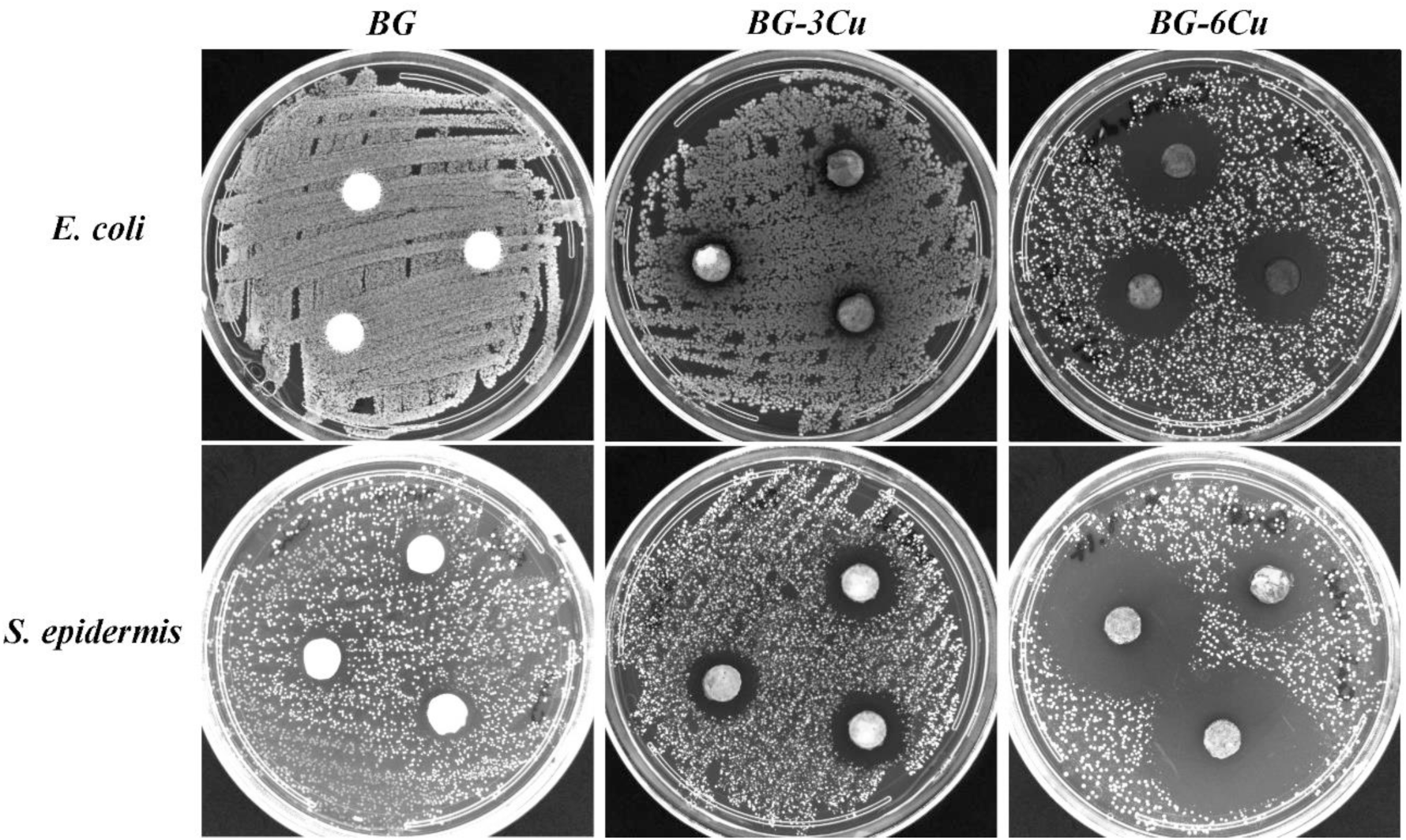
Antibacterial agar diffusion of *BG, BG-3Cu*, and *BG-6Cu* glasses against E. coli and S. epidermis.

## Conclusion

The substitution of CuO for SiO_2_ was investigated in order to study the structural effects and antibacterial efficacy in the bioactive glass compositions. Results from DTA, XPS, and NMR indicated that the copper ions act as network modifiers in the glass structure. Addition of copper resulted in a higher number of non-bridging oxygens at the expense of birding-oxygen. This resulted in a more disrupted network in copper-containing glasses, which in turn could increase the solubility and bioactivity of the glass in physiological solutions. The results from the agar diffusion antibacterial test indicated significantly higher efficacy of copper-containing glasses, as potential alternatives for antibiotics in the biomedical glasses.

## References

1. Jones, J. R., Acta Biomater., 2013, 9 (1), 4457–86. https://doi.org/10.1016/j.actbio.2012.08.023

2. Baino, F., Hamzehlou, S., Kargozar, S., J. Funct. Biomater., 2018, 9 (1), 25. https://doi.org/10.3390/jfb9010025

3. Rahaman, M. N., Day, D. E., Bal, B. S., Fu, Q., Jung, S. B., Bonewald, L. F., Tomsia, A. P., Acta Biomater., 2011, 7 (6), 2355–73. https://doi.org/10.1016/j.actbio.2011.03.016

4. Rabiee, S. M., Nazparvar, N., Azizian, M., Vashaee, D., Tayebi, L., Ceram. Int., 2015, 41 (6), 7241–7251. https://doi.org/10.1016/j.ceramint.2015.02.140

5. Hoppe, A., Guldal, N. S., Boccaccini, A. R., Biomaterials, 2011, 32 (11), 2757–74. https://doi.org/10.1016/j.biomaterials.2011.01.004

6. Ribas, R. G., Schatkoski, V. M., do Amaral Montanheiro, T. L., de Menezes, B. R. C., Stegemann, C., Leite, D. M. G., Thim, G. P., Ceram. Int., 2019, 45(17), 21051–21061. https://doi.org/10.1016/j.ceramint.2019.07.096

7. Chon, S. S., Piraino, L., Mokhtari, S., Krull, E. A., Coughlan, A., Gong, Y., Mellott, N. P., Keenan, T. J., Wren, A. W., J. Non-Cryst. Solids., 2018, 490, 1–12. https://doi.org/10.1016/j.jnoncrysol.2018.03.006

8. Balamurugan, A., Balossier, G., Laurent-Maquin, D., Pina, S., Rebelo, A. H. S., Faure, J., Ferreira, J. M. F., Dent. Mater., 2008, 24 (10), 1343–1351. https://doi.org/10.1016/j.dental.2008.02.015

9. Fiume, E., Barberi, J, Verne, E., Baino, F., J. Funct. Biomater., 2018, 9 (1). https://doi.org/10.3390/jfb9010024

10. Sanz-Herrera, J. A., Boccaccini, A. R., Int. J. Solids Struct., 2011, 48 (2), 257–268. https://doi.org/10.1016/j.ijsolstr.2010.09.025

11. Tilocca, A., Int. J. Mater. Chem., 2010, 20 (33), 6848–6858. https://doi.org/10.1039/C0JM01081B

12. Wang, Z., Cheng, L., J. Alloys Compd., 2014, 597, 167–174. https://doi.org/10.1016/j.jallcom.2014.01.232

13. Kaur, G., Pandey, O. P., Singh, K., Homa, D., Scott, B., Pickrell, G., J. Biomed. Mater. Res., Part A., 2014, 102 (1), 254–274. https://doi.org/10.1002/jbm.a.34690

14. Lázaro, G. S., Santos, S. C., Resende, C. X., dos Santos, E. A., J. Non-Cryst. Solids., 2014, 386, 19–28. https://doi.org/10.1016/j.jnoncrysol.2013.11.038

15. Babu, M. M., Prasad, P. S., Rao, P. V., Govindan, N. P., Singh, R. K., Kim, H.-W., Veeraiah, N., Ceram. Int., 2019, 45 (17), 23715–23727. https://doi.org/10.1016/j.ceramint.2019.08.087

16. Waltimo, T., Brunner, T. J., Vollenweider, M., Stark, W. J., Zehnder, M., J. Dent. Res., 2007, 86 (8), 754–7. https://doi.org/10.1177%2F154405910708600813

17. Mokhtari, S., Skelly, K. D., Krull, E. A., Coughlan, A., Mellott, N. P., Gong, Y., Borges, R., Wren, A. W., J. Mater. Sci., 2017, 52 (15), 8886–8903. https://doi.org/10.1007/s10853-017-0945-5

18. Finney, L., Vogt, S., Fukai, T., Glesne, D., Clin. Exp. Pharmacol. Physiol., 2009, 36 (1), 88–94. https://doi.org/10.1111/j.1440-1681.2008.04969.x

19. Mokhtari, S., Wren, A. W., Biomed. Glasses, 2019, 5 (1), 13–33. https://doi.org/10.1515/bglass-2019-0002

20. Li, Z., Khun, N. W., Tang, X.-Z., Liu, E., Khor, K. A., J. Mech. Behav. Biomed. Mater., 2017, 65, 77–89. https://doi.org/10.1016/j.jmbbm.2016.08.007

21. Brückner, R., Chun, H.-U., Goretzki, H., Sammet, M., J. Non-Cryst. Solids., 1980, 42 (1-3), 49–60. https://doi.org/10.1016/0022-3093(80)90007-1

22. Maekawa, H.; Maekawa, T.; Kawamura, K.; Yokokawa, T., J. Non-Cryst. Solids., 1991, 127 (1), 53–64. https://doi.org/10.1016/0022-3093(91)90400-Z

23. Angeli, F., Gaillard, M., Jollivet, P., Charpentier, T., Geochim. Cosmochim. Acta., 2006, 70 (10), 2577–2590. https://doi.org/10.1016/j.gca.2006.02.023

24. Stebbins, J. F., Nature, 1987, 330 (6147), 465. https://doi.org/10.1038/330465a0

25. Vu’o’ng, B. X., Hiêp, Đ. T., Glass Phys. Chem., 2016, 42 (2), 188–193. https://doi.org/10.1134/S108765961602005X

26. Gross, T. M., Lahiri, J., Golas, A., Luo, J., Verrier, F., Kurzejewski, J. L., Baker, D. E., Wang, J., Novak, P. F., Snyder, M. J., Nat. Commun., 2019, 10 (1), 1979. https://doi.org/10.1038/s41467-019-09946-9

